# Conformational Plasticity of the ClpAP AAA+ Protease Couples Protein Unfolding and Proteolysis

**DOI:** 10.1101/820209

**Authors:** Kyle E. Lopez, Alexandrea N. Rizo, Eric Tse, JiaBei Lin, Nathaniel W. Scull, Aye C. Thwin, Aaron L. Lucius, James Shorter, Daniel R. Southworth

**Author notes:** These authors contributed equally to this work (Kyle E. Lopez, Alexandrea N. Rizo). Correspondence to: D.R.S.

## Abstract

The ClpAP complex functions as a “bacterial proteasome” that simultaneously unfolds and degrades proteins targeted for destruction. ClpA utilizes two AAA+ domains per protomer to power substrate unfolding and translocation into the ClpP proteolytic chamber. To understand this mechanism, we determined high-resolution structures of wildtype *E. coli* ClpAP in distinct substrate-bound states. ClpA forms a spiral with substrate contacts across both AAA+ domains, while protomers at the seam undergo nucleotide-specific rearrangements indicating a conserved rotary mechanism. ClpA IGL loops extend flexibly to bind the planar, heptameric ClpP surface and support a large ClpA-P rotation that re-orients the translocation channel. The symmetry mismatch is maintained at the spiral seam through bind and release states of the IGL loops, which appear precisely coupled to substrate translocation. Thus, ClpA rotates around the apical surface of ClpP to processively translocate substrate into the protease.

## Main

The Hsp100 (Clp) family of proteins, widely present in bacteria and eukaryotes, function as protein unfoldases and disaggregases^1,2^. Some family members can assemble into large proteolytic machines homologous to the 26S proteasome and serve critical roles in targeted protein degradation and quality control^3–7^. Proteolysis requires substrate recognition and ATP hydrolysis-driven unfolding by a AAA+ Hsp100 complex, which unfolds and translocates the substrate into a proteolytic chamber^8–12^. The highly conserved serine protease, ClpP forms this chamber as a double ring of heptamers^13,14^ which partner with 1-2 ClpX or ClpA AAA+ hexamers in bacteria, assembling into single and double-capped complexes^15–17^. To promote client degradation, ClpXP and ClpAP are aided by SspB^18,19^ and ClpS^20,21^, specificity adaptors that promote recognition of substrates including those containing the ssrA degron^22,23^ and N-end rule substrates^24^, respectively. Other substrates, such as the RepA DNA-binding protein, recognized by ClpA, are remodeled or degraded in support of specific cellular functions^3,25^.

Hsp100 interactions with ClpP involve a hexamer-heptamer symmetry-mismatch, which is conserved among proteolytic machines that include the 20S core particle^3,6^. Contacts are mediated by IGF/L-motif loops in ClpX or ClpA and hydrophobic binding pockets on the apical surface of ClpP^6,26^. Engagement of these loops triggers an open-gate conformational change of adjacent N-terminal loops on ClpP, facilitating substrate transfer to proteolytic sites^27–29^. Indeed, the acyldepsipeptide class of antibiotics (ADEPs) compete for binding to these pockets and stabilize an open-gate conformation, thereby converting ClpP to an uncontrolled, general protease^30–33^. How these Hsp100-ClpP interactions are coordinated during active unfolding and translocation is unknown.

ClpA contains two nucleotide-binding AAA+ domains (D1 and D2) per protomer which power unfolding^34^. Structures of related disaggregases, Hsp104 and ClpB, identify the substrate-bound hexamer adopts a right-handed spiral in which conserved, Tyr-bearing pore loops across both AAA+ domains contact and stabilize the polypeptide substrate via backbone interactions spaced every two amino acids^35–38^. Distinct substrate-bound states reveal a ratchet-like mechanism defined by the spiral arrangement, in which the ATP hydrolysis cycle drives substrate release at the lower position and re-binding to the topmost position along the substrate^1,36^. A similar spiral architecture and array of substrate contacts has now been identified for many AAA+ machines, supporting a universal rotary translocation mechanism^39–42^. However, it is unclear how this mechanism is coupled to proteolysis, or how interactions are maintained with the planar, heptameric surface of the protease ring during translocation cycling.

Here, we sought to determine the structural basis for coupled protein unfolding and proteolysis by the ClpAP complex. Using the slowly hydrolysable ATP analog, ATPγS, and a RepA-tagged GFP substrate, we determined cryo-EM structures of intact, wildtype ClpAP from *E. coli* to ~3.0 Å resolution stalled in distinct substrate translocation states. Two states of the hexamer-heptamer interface are identified which reveal the IGL loops undergo release and rebinding at the ClpA spiral seam and support a rotation between ClpA and ClpP that re-positions the substrate channel. These changes in the IGL loops coincide with nucleotide-specific rearrangements in the AAA+ domains that advance substrate interactions, supporting a stepwise rotary translocation cycle. Thus, these structures reveal a long-range conformational network that enables ClpA contacts with the planar surface of ClpP to be precisely coordinated with the conserved ratchet mechanism, thereby enabling ATP-powered substrate unfolding and transfer into the ClpP chamber where substrate is degraded.

## Results

### Architecture of the Substrate-Bound ClpAP Complex

To determine structures of a substrate-bound ClpAP complex, a RepA-GFP substrate was tested containing the first 25 residues of RepA (RepA^1–25^-GFP), and includes sequences established as sufficient for ClpA recognition and degradation ^10,43^. RepA-GFP substrates are actively proteolyzed by ClpAP and previously established for monitoring unfolding by ClpA^43,44^. Incubations included the slowly hydrolysable ATP analog, ATPγS, in order to improve binding and slow or stall translocation (Supplementary Fig. 1a-b). By size exclusion chromatography (SEC), ClpAP forms a stable supercomplex and binds to GFP containing RepA^1–25^ after incubation with ATPγS. The high molecular weight peak fractions containing ClpAP-RepA^1–25^-GFP complex were collected for subsequent cryo-EM analysis (Supplementary Fig. 1a). ClpAP proteolysis of RepA^1–25^-GFP was assessed in the presence of saturating ATP or ATPγS by SDS-PAGE (Supplementary Fig. 1b), revealing near complete degradation within 30 minutes with ATP and no apparent degradation with ATPγS (Supplementary Fig. 1b). Notably, a predominant RepA^1–25^-GFP cleavage product is observed at shorter times indicating the RepA tag is likely rapidly degraded prior to unfolding and proteolysis of GFP (Supplementary Fig. 1b). Together these results indicate that ClpAP actively proteolyzes RepA^1–25^-GFP with ATP and forms a stable complex with RepA^1–25^-GFP in the presence of ATPγS, likely becoming stalled in a substrate bound state.

In reference-free 2D class averages, side views of ClpP particles double-capped with ClpA predominate and in top-views the density indicates the presence of two rings of different diameters, indicative of ClpAP (Fig. 1a and Supplementary Fig. 1c). While two ClpA hexamers were identified, one typically showed well-resolved features compared to the other, indicating preferred alignment likely due to flexibility across the double-capped complex. Separate 3D classifications of two independent datasets were performed for the double-capped complex and both yielded two distinct conformations (Supplementary Fig. 1j). As with 2D analysis, one ClpA hexamer showed improved features over the other. The final maps refined to an overall resolution of 3.1 and 3.0 Å for the two states, herein after referred to as the Engaged (ClpAP^Eng^) and Disengaged (ClpAP^Dis^) states, respectively, based on the binding states of the IGL loops (Supplementary Fig. 1d-h). In both states the D1 and D2 AAA+ rings of the ClpA hexamer adopt a right-handed spiral with the D2 ring contacting the planar, heptameric surface of ClpP via the IGL loops (Fig. 1b). ClpA is comprised of protomers P1-P6 with P1 at the lowest and P5 at the highest position of the spiral, while P6 defines the seam interface (Fig. 1b). This architecture is similar to related ClpB and Hsp104 double-ring disaggregases in the substrate-bound states ^35–37^. Resolution is the highest for ClpP (~2.5 Å), while ClpA is more variable (~2.5-4 Å for ClpAP^Dis^ and ~3-6 Å for ClpAP^Eng^), with the spiral seam protomers (P1, P5 and P6) at lower resolutions due to their flexibility (Supplementary Fig. 1e-f). The high-resolution of the maps permitted accurate atomic models to be built for the full ClpAP complex (Fig. 1c, Supplementary Fig.1k-m; Table 1). Density for the flexible N-terminal (NT) domain of ClpA (residues 1-168) was not well-resolved, and thus was not modeled.

**Fig. 1:**
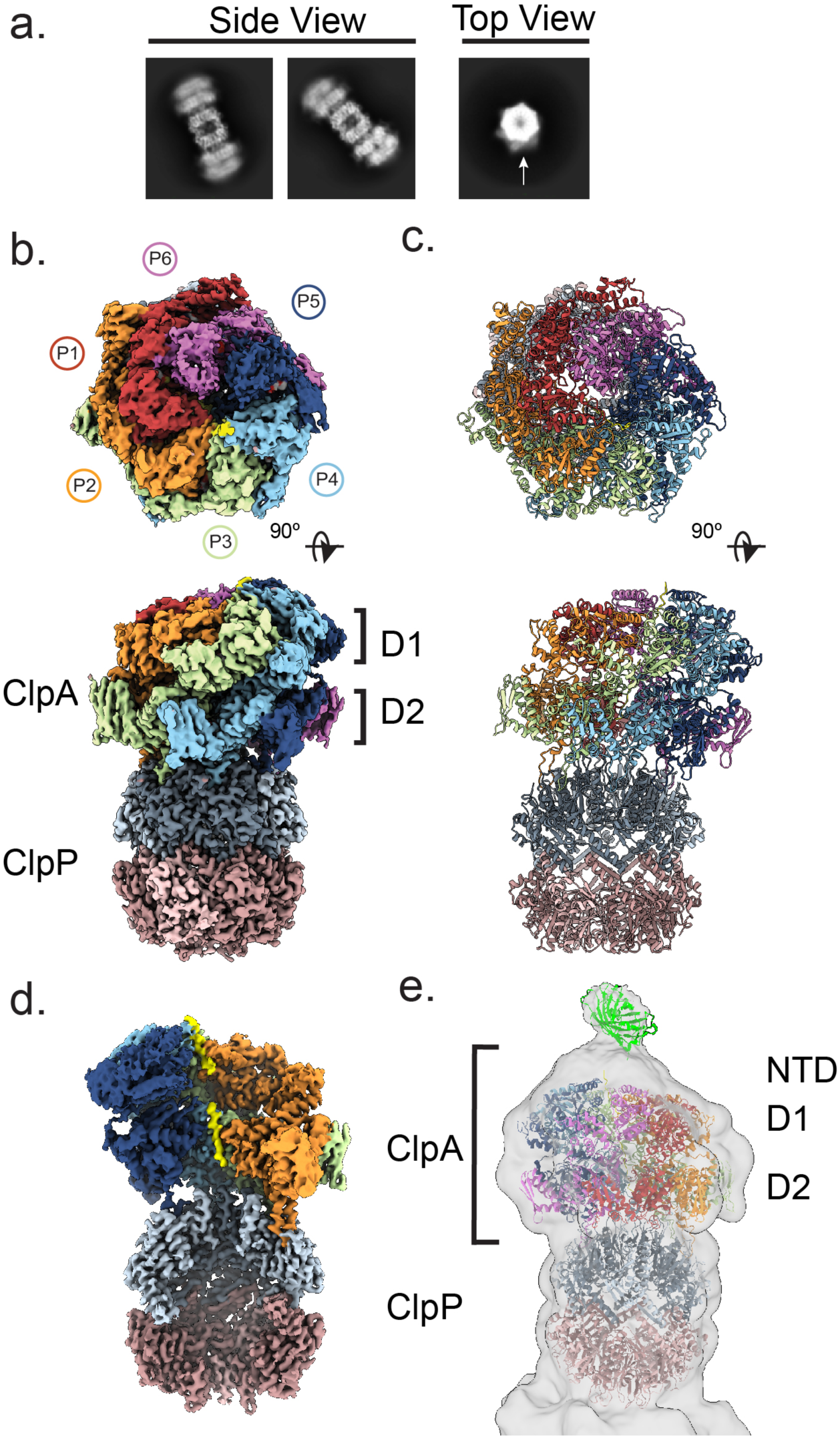
Cryo-EM structure of the substrate-bound ClpAP complex. **a,** 2D class averages of double-capped ClpAP. Both ClpA (arrow) and ClpP rings identified in top views. **b,** Top and side views of the final map and **c**, model of ClpAP^Eng^. ClpA is colored by individual protomer, as indicated. **d,** Channel view showing substrate peptide bound to ClpA (yellow). **e,** Low-pass filtered map showing globular density docked with GFP (pdb:1GFL) and additional N-terminal ClpA density (NTD)

Density corresponding to an unfolded polypeptide substrate is well-resolved along the ClpA channel for the D1 and D2 domains and was initially modeled as 10 and 11-residue poly-Ala chains respectively (Fig. 1d). Continuous density along the channel is identified for the substrate at reduced thresholds, indicating a single polypeptide chain is bound (Supplementary Fig. 1i). However, the connecting density between domains, corresponding to 5 residues based on the ClpB and Hsp104 structures ^35–37^, is poorly resolved and was not modeled. Notably, in low-passed filtered maps of the final reconstruction, globular density at the entrance to the ClpA channel is visible at a reduced threshold that approximately corresponds to a GFP molecule (Fig. 1e). These data along with the SEC analysis of the RepA-GFP constructs and proteolysis data indicate that ClpAP under these conditions with ATPγS is likely stalled with the 25-residue RepA sequence in the ClpA channel while GFP may remain folded at the apical surface.

### Large ClpA-P Rotation Coincides with IGL Loop Bind and Release

As noted above, two distinct conformations of substrate-bound ClpAP refined to high resolution. In the Engaged state, IGL loops from all 6 ClpA protomers are bound to pockets in ClpP with the remaining empty pocket of the heptamer positioned beneath the ClpA spiral seam between protomers P1 and P6 (Fig. 2a, 2c). Surprisingly, in the Disengaged state the IGL loop of protomer P1, which is at the lowest position along the substrate, is released from the ClpP pocket, resulting in two neighboring empty pockets at the ClpA seam (Fig. 2b, 2d). No conformational differences are identified for ClpP between the two states (RMSD =0.65). However, upon alignment of the ClpP portion of structures, the entire ClpA hexamer is identified to shift position through a pivot at the hexamer-heptamer interface (Fig. 2a-b, Supplementary Video 1). In the Engaged state the ClpA channel is offset from the ClpP open gate and central chamber by ~9°, while in the Disengaged state the channel shifts substantially to become offset by ~17° from ClpP. In a morph between these two states, a rotation of ClpA is visualized in which protomers P4-P6 tilt towards ClpP while P1-P3 tilt away. These changes coincide with disengagement of the P1 IGL loop, indicating loss of this binding interaction may facilitate rotation of ClpA.

**Fig. 2:**
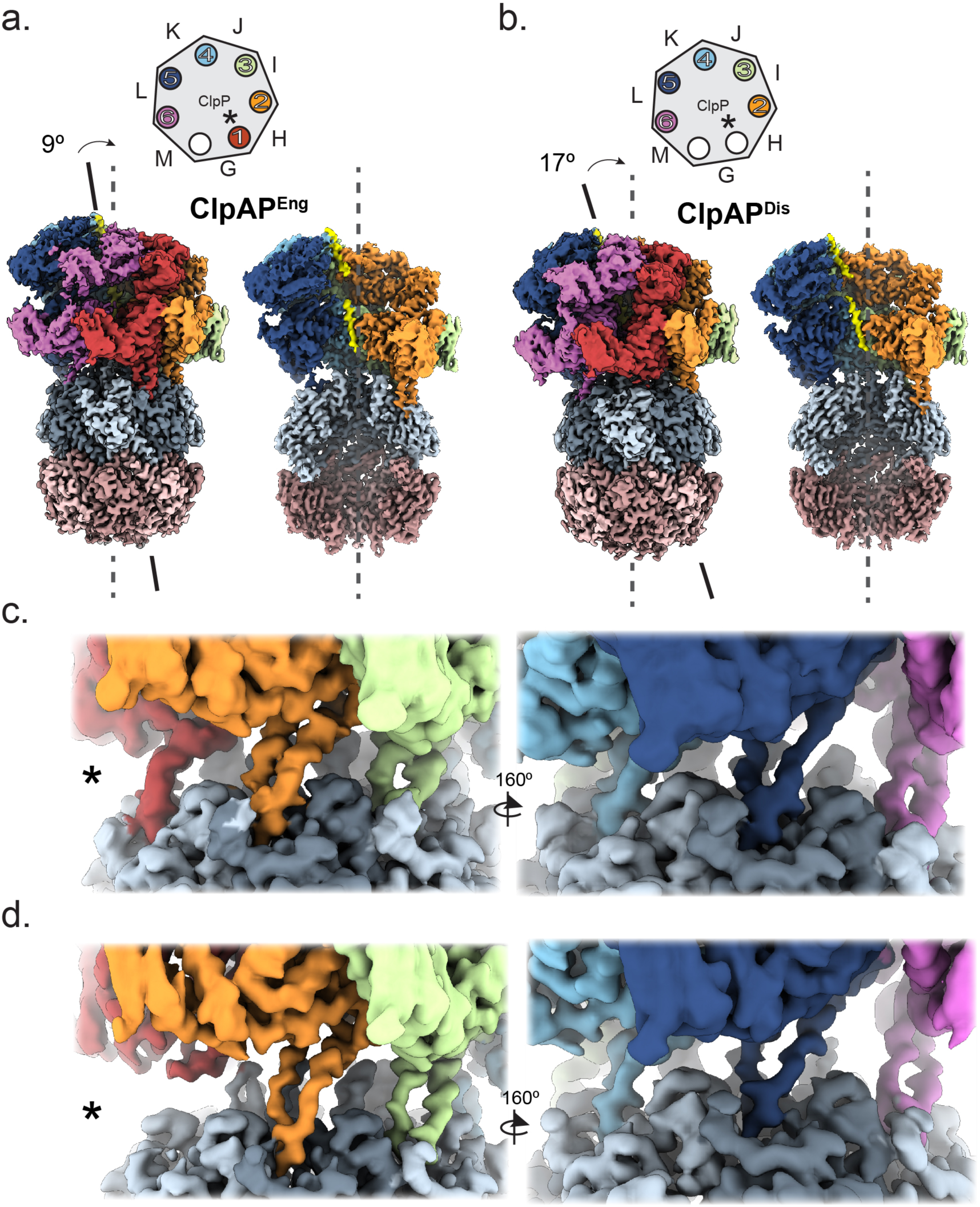
IGL loop engaged and disengaged states of ClpAP. **a,** ClpAP^Eng^ and **b,** ClpAP^Dis^ cryo-EM maps showing degree offset (arrow) of the ClpA channel axis (solid line) and substrate position (yellow density) compared to the ClpP pore and proteolytic chamber (dashed line). Schematic (above) shows occupancy of the ClpA IGL-loops (circles, colored and numbered by protomer) around the ClpA heptamer, with the empty IGL pockets (white circles) and ClpA protomers indicated (letters) for the two states. **c,** Cryo-EM density of the ClpA-P interface showing IGL loop binding to ClpP in the ClpAP^Eng^ and **d**, ClpAP^Dis^ States. The IGL loop from protomer P1 which disengages ClpP is indicated (*).

While ClpAP^Eng^ and ClpAP^Dis^ were the two predominant states that refined to high resolution, comparison of the different classes of the double-capped ClpAP complex at lower-resolution revealed additional ClpA-P orientations, indicating that the arrangement is dynamic, with ClpA appearing to rotate about the hexamer-heptamer interface (Supplementary Fig. 2a-b). Resolution of these regions was not sufficient to define the IGL loop organization but likely involves release from ClpP pockets. Overall, we identify that the ClpA position on the ClpP interface is highly dynamic, likely reflecting the conformational changes in ClpA required for substrate translocation. Based on the ClpAP^Eng^ and ClpAP^Dis^ states, the ClpA rotations are supported by precise bind and release events of the IGL loop of the P1 protomer positioned at the spiral seam.

### IGL Loop Plasticity Enables ClpP Engagement by the ClpA Spiral

Previous crystal structures of ClpA were unable to resolve the IGL loops due to flexibility, but biochemical data for ClpX IGF loops suggest that they make static interactions with ClpP and all 6 IGF loops are required for optimal activity^45^. In the ClpAP^Eng^ and ClpAP^Dis^ structures, density for the ClpA IGL loops is well-defined enabling precise modeling for all the loops that are engaged in the ClpP pockets (Supplementary Fig. 3a-b). The IGL loop region extends from residues N606 and T637 in the base of the D2 large subdomain as two short α helices. Residues 616-620 form the flexible loop, which extends into the hydrophobic binding pocket on ClpP, resulting in ~600 Å^2^ of buried surface area compared to the empty pocket (Fig. 3a, left). The IGL loop binding pocket is formed by the interface of two ClpP protomers and includes α helices B and C from one protomer and a 3-strand β sheet (strands 1, 2, and 3) and the C-terminal (CT) strand from the adjacent protomer (Fig. 3a). The loop residues I617, G618, L619, and I620 bind a hydrophobic region in the pocket comprised of A52, L48, F49, and F82 in α helices of one protomer and L23, Y60, Y62, I90, M92, F112, L114, L189 in the adjacent protomer (Fig. 3a, middle). Additional electrostatic contacts likely stabilize the loop as well, including R192 in the CT strand, and E26, which appear to interact with H621 and R614 and Q622, respectively (Fig. 3a, right). There are also a few hydrogen bonds between the IGL loop, L619 and the adjacent ClpP protomers, R192 and Y62 as well as within the IGL loop itself S616 and H621.

**Fig. 3:**
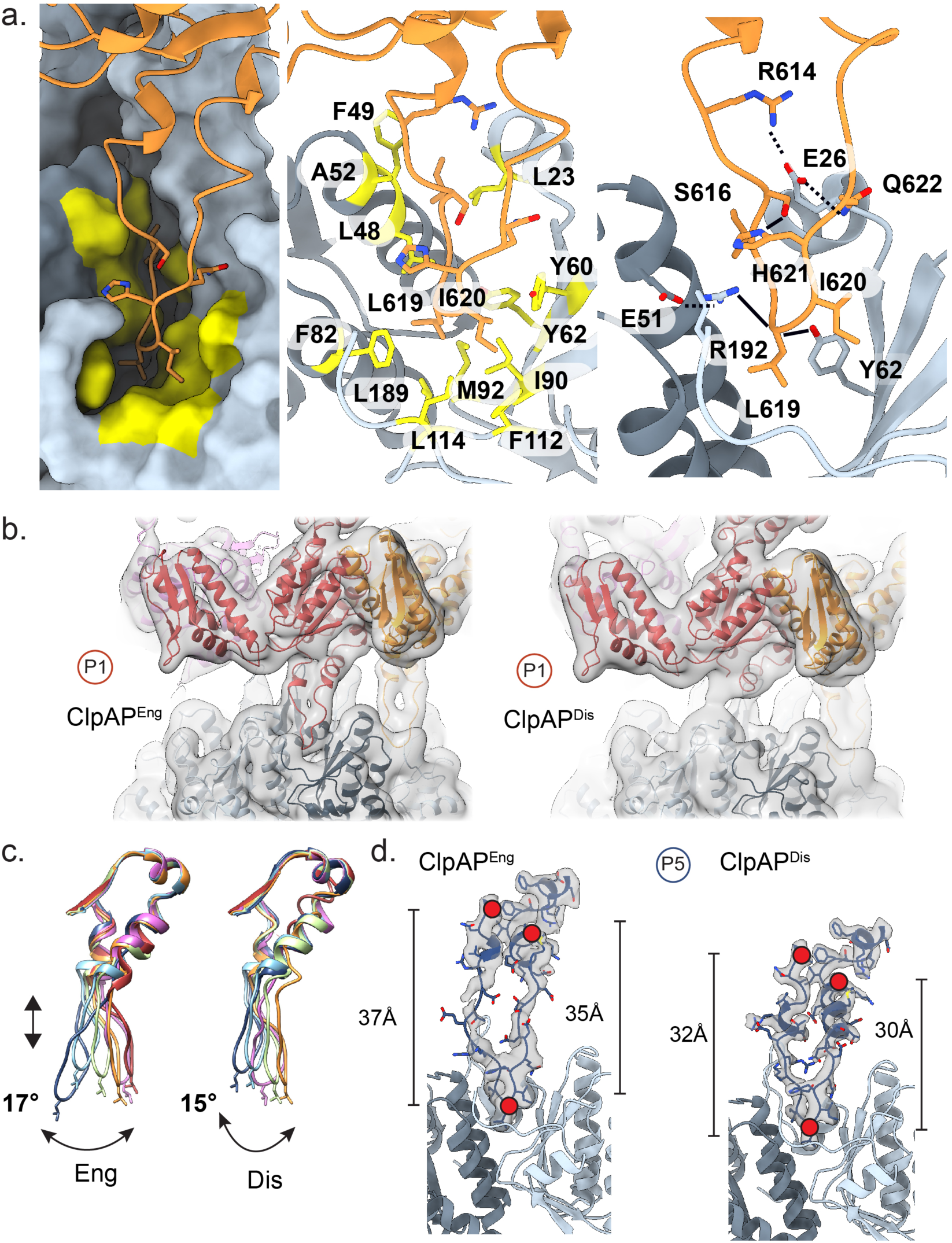
IGL loop interactions and conformational flexibility. **a,** Representative view of a bound IGL loop (orange, ribbon view) positioned in the binding pocket of ClpP shown in surface view with hydrophobic residues colored in yellow (left), and shown in ribbon views with hydrophobic interactions (middle) and electrostatic contacts (right) labeled. **b,** Low-pass filtered map and model showing P1 IGL-loop density extends into the IGL pocket for ClpAP^Eng^ (left) and is disengaged for ClpAP^Dis^, contacting the adjacent apical ClpP surface (right). **c,** Overlay of IGL loops (colored by protomer) of ClpAP^Eng^ (left) and ClpAP^Dis^ (right). IGL loops are aligned to connecting residues 638-649. **d,** Map and model of the P5 IGL-loop for ClpAP^Eng^ (left) and ClpAP^Dis^ (right) showing extended and compact conformations, respectively, based on distance measurements between loop residues 605-619 and 633-619 (red dots).

Similar to the other loops, the P1 IGL loop is well-resolved in the Engaged state and bound in the ClpP pocket (Fig. 3b, left). However, for ClpA^Dis^, loop residues 609-624 appear disordered and could not be modeled. Notably, in a lower threshold, filtered map the ClpP pocket remains empty while density extends laterally from the loop helices to contact the apical surface of ClpP (Fig. 3b, right). Thus, for ClpA^Dis^ the IGL loop is shifted clockwise and becomes positioned adjacent the surface between the two empty binding pockets. This shift is due primarily to the large rotation of the ClpA hexamer relative to ClpP between the Engaged and Disengaged states. (Supplementary Video 1).

When the top of the loops (residues 638-649) are aligned in each protomer of ClpA, substantial rearrangements are identified in both states, revealing the conformational plasticity of the loops around the hexamer (Fig. 3c). In general, the loops rotate around residues (613 and 624) at the end of the two short helices, with the most significant changes occurring between the adjacent P5 and P6 loops, which rotate by ~17°. This rotation is notable because P6 is at the spiral seam and unbound to substrate while P5 is bound at the highest position of the substrate, similar to other AAA+ structures (discussed below). This protomer pair are the most conformationally distinct based on their position along the channel axis, illustrating how the large flexibility of the IGL loops facilitates engagement of ClpP despite the variable position of the ClpA protomers due to their spiral arrangement and symmetry mismatch with ClpP. The IGL loops additionally undergo expansion and contraction at different positions around the hexamer and between the Engaged and Disengaged states (Fig. 3c, and Supplementary Fig. 3c). This is most notable for protomer P5, which extends by 37 and 35 Å in the Engaged state compared to 32 and 30 Å in the Disengaged state, thereby supporting the large pivot of ClpA across ClpP (Fig. 3d). Remarkably, the extension of the IGL loops occurs through a partial unfolding of both connecting helices (residues 609-613 and 614-629), which could additionally serve to support the variable positions of the ClpA spiral relative to ClpP.

### ClpP Structure and N-terminal Gating

The flexible N-terminal loop residues of ClpP (1-18) form a pore on the apical surface that functions as a substrate gate, which is allosterically controlled by engagement of the adjacent IGF/L binding pockets by ClpX/A or ADEP compounds ^33^. In both ClpAP^Eng^ and ClpAP^Dis^ structures, the ClpP NT loops from each protomer are well-resolved and adopt an extended configuration resulting in an open gate conformation that is positioned adjacent the ClpA translocation channel, ~30 Å away from where substrate is resolved (Fig. 4a and Supplementary Fig. 4c-d). This is distinct from crystal structures showing the NT loops adopt an asymmetric open-gate arrangement ^46^, but similar to ADEP-bound structures where all the loops are in an extended conformation ^31,33^. Additionally, no contact is observed between the NT loops and ClpA (Fig. 4a), which may be distinct compared to ClpXP, in which NT loops have been identified to contact the ClpX pore-2 loops ^45^.

**Fig. 4:**
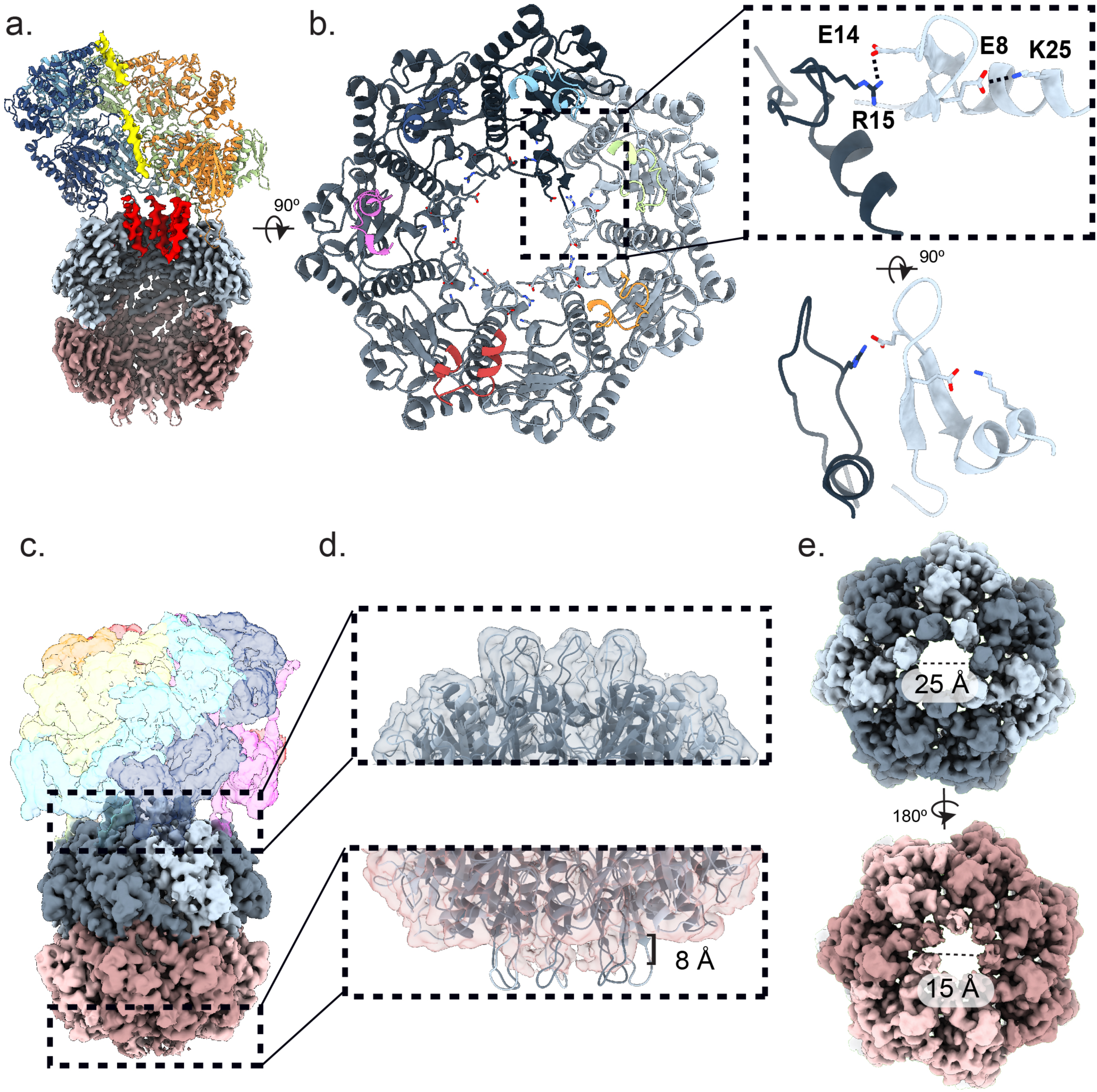
Structure of ClpP and NT gating loops. **a,** Channel view of ClpAP^Eng^ highlighting the ClpP NT gating loops (red). **b,** (left) Top view of ClpP NT loops with ClpA IGL loops shown, colored by protomer and ClpA IGL loops displayed; and (right) top of NT loop pair with cis (E8-K25) and trans (R15-E14) salt-bridge contacts. **c,** Side-view map of single-capped ClpAP complex. **d,** Expanded map+model view of ClpP NT loop region for the ClpA-bound, open-gate and unbound closed-gate conformations, with distances shown. **e,** Top views showing ClpP pore diameter for the (top) open- and (bottom) closed-gate conformations.

We identify two specific interactions: one across the ClpP NT loops and one with an adjacent helix A in the IGF/L pocket, which have not been previously characterized and appear to stabilize the open gate conformation (Fig. 4b). A salt-bridge contact between residues R15 in one loop and E14 in the clockwise loop is identified in each protomer (Fig. 4b and Supplementary Fig. 4e). Additionally, a potential salt-bridge contact involving E8 and K25 is also observed which may additionally stabilize the loop orientation (Fig. 4b and Supplementary Fig. 4f). Notably, K25 is located in a helix that comprises part of the hydrophobic, IGL-binding pocket (Fig. 4b). Thus, this interaction may be involved in the allosteric gating mechanism.

In an initial ClpAP dataset, we identified a population of single-capped complexes which resolved into one 3D class (Fig. 4c and Supplementary Fig. 4a-b), enabling us to characterize the open and closed-gate conformations in one structure. While the resolution of the NT loops was not sufficient to model the closed conformation, at lower threshold values, density for the loops on the unbound end of ClpP appears to extend ~8 Å from ClpP, while density for the ClpA-bound end NT loops extends ~16 Å (Fig. 4d). Additionally, the pore diameter is identified to be ~25 Å for the ClpA-bound end of ClpP, which is substantially wider compared to the unbound end, which is ~15 Å (Fig. 4e). Thus, we identify the NT loop gating mechanism is specifically triggered by engagement of the cis-bound ClpA IGL loops which may allosterically regulate the NT loops through salt bridge contacts that stabilize the extended loop arrangement.

### ClpA Substrate Contacts and Translocation States

To improve the resolution of the ClpA pore loop interactions and the seam protomers, particle subtraction and focused refinement of the ClpA hexamer was performed (Supplementary Fig. 5a). This resulted in an overall estimated resolution of 3.2Å and 3.1Å for the ClpA^Eng^ and ClpA^Dis^ focused maps, respectively (Supplementary Fig. 5a,e). Some improvement in the map density for the seam protomers and substrate contacts was observed, therefore models were further refined using these maps to characterize the substrate interactions (Supplementary Fig. 5b-d,g). Similar to other AAA+ structures, the conserved Tyr-pore loops in the D1 and D2 of ClpA extend into the channel and form a spiral of substrate interactions spaced every two amino acids along the polypeptide (Fig. 5a). The D1 stabilizes a 9-residue segment through direct contact by Y259, which intercalates between the substrate side chains and contacts the backbone (Fig. 5b). The conserved flanking residues, K258 and R260, extend laterally to make electrostatic contacts with the upper and lower adjacent pore loops (D262 and E264), similar to what is identified for ClpB D1^37,38^ (Supplementary Fig. 5f). In the ClpAP^Eng^ structure 4 protomers, P1-P4, contact the substrate while protomers P5 and P6 are unbound, with Y259 positioned ~17 Å and ~20 Å away from the substrate, respectively (Fig. 5b and Supplementary Video 2). Upon comparing the ClpAP^Eng^ and ClpAP^Dis^ structures, the substrate bound protomers, P1-P4, show no conformational changes. However, in the ClpAP^Dis^ structure both P5 and P6 pore loops shift closer to the polypeptide substrate by 8 and 4 Å respectively, and move up the channel axis (Fig. 5b and Supplementary Video 2). Notably for ClpAP^Dis^, P5 Y259 becomes positioned adjacent the next substrate contact site, two residues above the P4 Y259 position, indicating a translocation step likely occurs upon transitioning from the Engaged to Disengaged states.

**Fig. 5:**
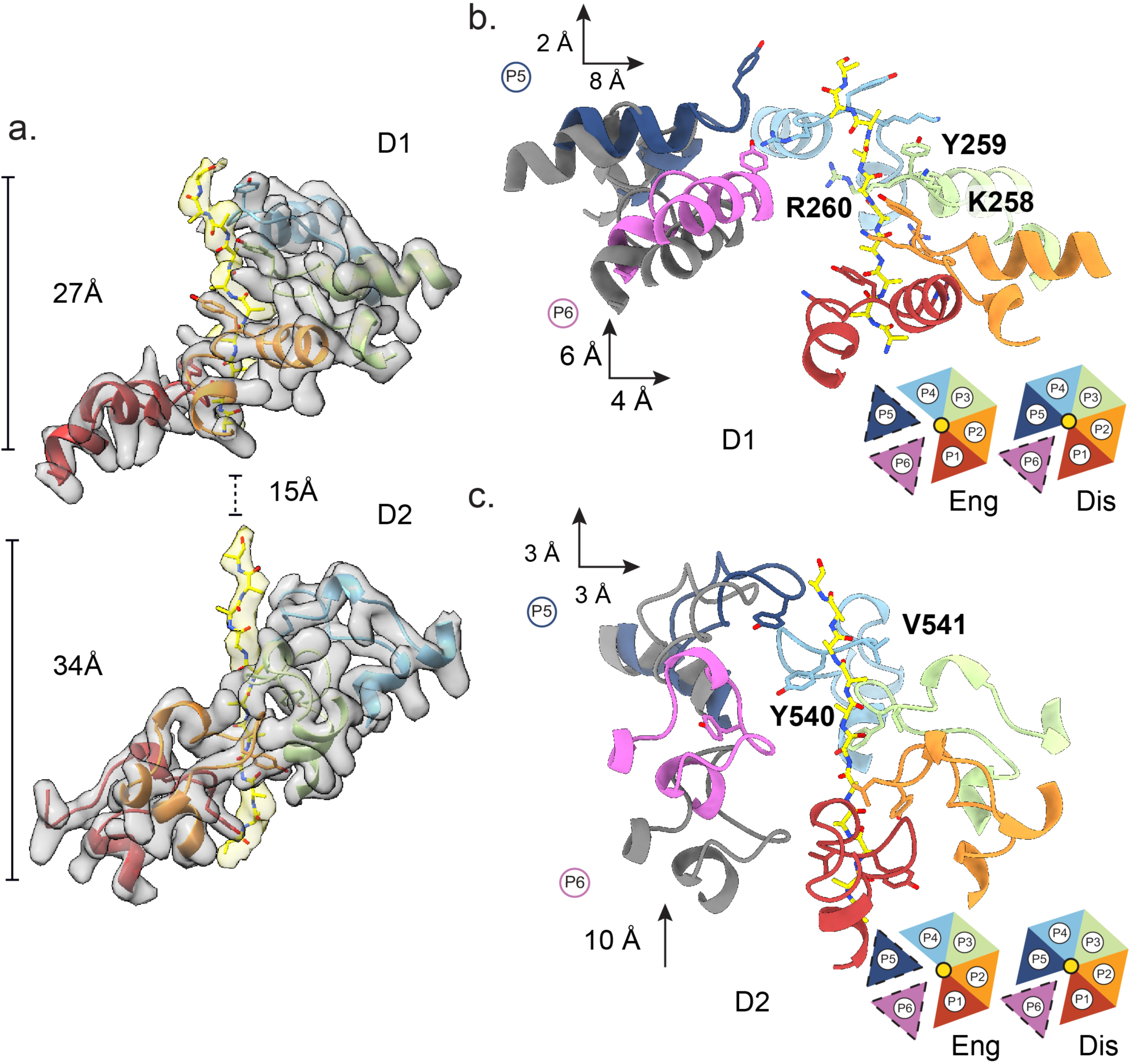
ClpA pore loop-substrate contacts and translocation states. **a,** Segmented map+model of the substrate-bound P1-P4 pore loops, colored by protomer, with substrate (yellow) for ClpAP^Eng^. Distances shown for the spacing of the D1 and D2 substrate segments. **b-c,** Model of the **b,** D1 and **c,** D2 pore loops and substrate for ClpAP^Eng^ (grey) and ClpAP^Dis^ (colored by protomer) with substrate-contacting residues shown. Distance shifts for the seam protomers, P5 and P6 between the engaged (grey) disengaged (blue and magenta) states. Schematic shows top view of pore loop (triangles)-substrate (yellow) contact arrangement for the indicated states.

The D2 similarly shows well-defined pore loop-substrate contacts for protomers P1-P4 in both states (Fig. 5c). These interactions stabilize a longer, 11 residue polypeptide segment and are primarily mediated by Y540 and V541, which form a clamp around the substrate backbone. As with the D1, the pore loops for P5 and P6 are unbound to substrate in the Engaged state, but shift up and toward the polypeptide in the Disengaged state (Fig. 5c and Supplementary Video 2). Notably, with this conformational change the P6 loop shifts a substantial, 10 Å along the substrate axis while the P5 loop rotates inward and contacts the substrate two residues above the D2 P4 position. Additional, pore-2 loops^47,48^ are present in both the D1 (residues: 292-302) and D2 (residues: 613-625), are conserved in ClpB and Hsp104 ^36–38^ and appear to make additional contributions to stabilizing the polypeptide in the channel. Within the pore-2 loops, residues A295 and A296 in the P4 D1 and H528 in the P3 and P4 D2 are positioned adjacent the substrate (Supplementary Fig. 5h). Considering the D2 pore-2 loops are adjacent the ClpP N-terminal gate (~10 Å for P1), substrate interactions by these loops may facilitate transfer to the ClpP chamber.

### Hydrolysis Drives Translocation and Conformational Coupling with ClpA

Similar to Hsp104 and ClpB, ATP hydrolysis activity in D1 and D2 is required for ClpA function ^49^. Both ClpA^Eng^ and ClpA^Dis^ structures show well resolved nucleotide pockets, therefore the nucleotide state of each NBD was assessed based on the ClpA focus map density for nucleotide and the position of the trans-activating Arg-finger residues (R339-R340 in the D1 and R643 in D2) (Supplementary Fig. 6a). For both D1 and D2 of ClpA^Eng^ and ClpA^Dis^ structures the substrate bound protomers P2-P4 are bound to ATPγS and the corresponding Arg-finger residues are adjacent the γ-phosphate, indicating an ATP, active state configuration (Fig. 6a and Supplementary Fig. 6a). Protomer P1, which contacts substrate at the lowest position in the channel appears bound to ADP and likely in a post-hydrolysis state for both the D1 and D2 in the ClpAP^Eng^ and ClpAP^Dis^ structures. Similarly, the clockwise P6 protomer, which is at the spiral seam and unbound to substrate also appears to be in an ADP-state for both D1 and D2. Although, R339-R340 and R643 from P5 are adjacent the nucleotide pockets, density for the sidechains was not well-resolved potentially due to the absence of the interacting γ-phosphate (Supplementary Fig. 6a). The clockwise protomer, P5, transitions from being unbound to substrate in ClpAP^Eng^ to bound at the uppermost position in ClpAP^Dis^ (Fig. 5b) but appears to be in an ADP state for the D1 and an ATP state for the D2 in both states. These nucleotide states are consistent with those observed for ClpB and Hsp104 as well as other substrate-bound AAA+ structures, supporting a clockwise rotary hydrolysis cycle. In this model, hydrolysis occurs at the spiral interface, likely initiating in protomer P6 at the lowest substrate contact position and thereby triggering substrate release. ATP re-binding stabilizes the substrate-bound state of the pore loops which we propose to occur at the protomer P5 position, enabling a two amino acid translocation step to occur while protomers P1-P4 remain bound in the same position. While larger step sizes have been measured for ClpA^50^, a 6.5 Å, two amino acid step is consistent with previous Hsp100 structures^35–38^ and kinetic experiments comparing ClpA and ClpAP^51^

**Fig. 6:**
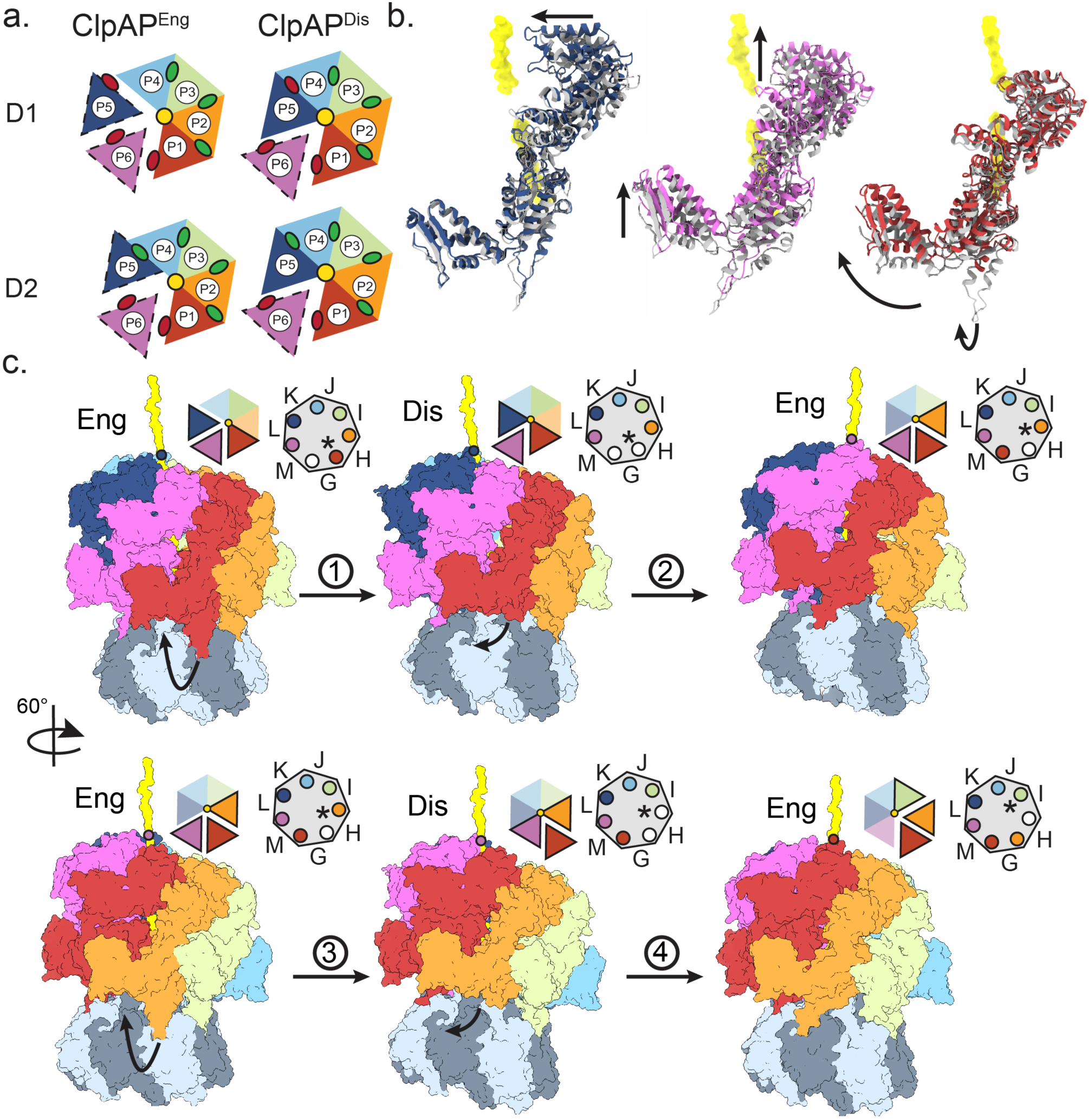
Nucleotide states and rotary translocation model for coupled unfolding and proteolysis. **a,** Schematic showing nucleotide states for the D1 and D2 of ClpAP^Eng^ (left) and ClpAP^Dis^ (right), determined based on difference maps (Supplementary Fig. 6). Protomer nucleotide states are denoted by colored circles (green for ATP and red for ADP). **b,** Overlay of the seam protomers P5 (left), P1 (middle), and P6 (right) for the ClpAP^Eng^ (grey) and ClpAP^Dis^ (colored) showing conformational shifts (arrows) supporting translocation step. **c,** Model for ClpAP processive substrate translocation cycle. Two translocation steps are depicted and coupled to IGL-loop release (step 1and 3) and binding the next clockwise IGL pocket on ClpP (step 2 and 4), indicated by arrows. Top view schematics show rotary cycle of substrate binding by ClpA (left) and occupancy ClpP IGL pockets. Rotating seam protomers are indicated in the hexagon by bold outline and the protomer at the lowest substrate contact site, which releases the IGL loop is indicated (*).

Together the P1, P5 and P6 seam protomers undergo large conformational changes between the Engaged and Disengaged states that result a concerted shift upward and toward the channel axis by the three protomers (Fig. 6b and Supplementary Video 3). Considering the P5 pore loops rotate towards the next substrate contact site, these changes also support a substrate translocation step involving a clockwise rotary mechanism of unfolding. Remarkably, these changes appear to be coupled to the P1 IGL loop interaction with ClpP. With the release of ClpP by P1 in the Disengaged state, the D2s of P6 and P1 together shift upward ~8 Å (Fig. 6b). Importantly, the P1-P6 interprotomer contacts are maintained at the D1 and D2 interfaces, enabling theses conformational changes to propagate across the length of protomers towards the P5 D1 and channel entrance, potentially facilitating binding to the next substrate contact site. Thus, binding and release of ClpP by the IGL loop appears to be precisely coupled to long-range conformational changes that drive substrate translocation, revealing the structural basis for how ClpP interaction can allosterically regulate unfolding^51,52^.

## Discussion

Structures of related substrate-bound AAA+ translocases, including the double-ring disaggregases ClpB and Hsp104, have revealed a spiral array of pore loop-substrate contacts and a dynamic translocation mechanism involving ATP hydrolysis-driven substrate release and rebinding at the seam interface of the hexamer^36–38,53^. To understand how these large conformational changes of the AAA+ hexamer could operate with an attached heptameric protease during coupled protein unfolding and proteolysis, we determined structures of the wildtype *E. coli* ClpAP complex bound to a RepA-tagged GFP substrate. Two distinct structures, ClpAP^Eng^ and ClpAP^Dis^, were determined and reveal the ClpA architecture and substrate interactions by the D1 and D2 pore loops are similar to the ClpB and Hsp104 homologs, supporting a conserved rotary mechanism for substrate translocation. Remarkably, comparison of these structures reveals how large rotations across the ClpA-P interface and plasticity of the ClpA IGL loops, including release from ClpP by the P1 protomer, could enable processive unfolding and proteolysis.

Based on the ClpAP^Eng^ and ClpAP^Dis^ structures, we propose a model for ClpAP function in which hexamer-heptamer symmetry mismatch is maintained with one empty IGL pocket of ClpP aligned beneath the spiral seam of ClpA. Meanwhile, the IGL loop of the adjacent protomer (P1) at the lowest substrate contact site, disengages from ClpP and re-binds to that clockwise empty pocket in a manner that is regulated by ATP hydrolysis and the conformational changes at the spiral seam that drive substrate translocation (Fig. 6c and Supplementary Video 4). ClpAP is proposed to operate processively, similar to other AAA+ proteolytic machines^2,54,55^, indicating that ClpA remains bound to ClpP during multiple cycles of ATP hydrolysis-driven unfolding. For a processive cycle, we propose that following the Engaged to Disengaged change (step 1), the IGL loop would next bind the clockwise pocket on ClpP (step 2), which is unbound in the previous step, thereby transitioning back to the Engaged state, but shifting the IGL position by one protomer on ClpP. The position of the spiral seam would additionally shift counterclockwise by one protomer, thus enabling the next substrate translocation step to occur with the change to the Disengaged state (step 3). Based on this model, processive translocation would therefore involve sequential clockwise shifts in the IGL binding site such that the unbound IGL pockets remain aligned with the seam position. Additionally, for every full cycle around the hexamer involving six translocation steps, ClpA will shift by one protomer position on ClpP. While other processive cycles are possible, this model is consistent with mapping the two structures we determined in a conformational morph around the hexamer (Supplementary Video 4), and maintains the spiral array of substrate contacts and position of the hexamer-heptamer mismatch. Additionally, this model agrees with rotary translocation models that we and others have proposed for AAA+ translocases, further revealing how ATP hydrolysis additionally powers allosteric changes at the ClpA-P interface in addition to substrate translocation, enabling substrate translocation to be coupled with transfer to ClpP and proteolysis.

While the IGL loop interactions with the ClpP hydrophobic pockets are identical at all positions, substantial flexibility occurs with the helices that connect the loops to the D2 base of ClpA. This plasticity is likely critical for ClpP engagement, given the spiral arrangement of the ClpA hexamer and thus variable distance to the apical surface of ClpP. Additionally, this flexibility supports the ratcheting conformations of the seam protomers during translocation and the large rotation of ClpA that occurs between the engaged and disengaged states (Fig. 2a,b). Notably, the loops show greater flexibility in the engaged state, including the large (~15°) rotation between protomers P5 and P6, and the extension and unfolding of the P5 IGL loop helices (Fig. 3c,d). This larger flex of the IGL loops, in particular at the spiral seam, may provide energetic constraints in the engaged state which could thereby facilitate release of the P1 IGL loop.

The nucleotide states of ClpA are similar to the arrangement identified in related AAA+ complexes^35–37,42^ in which the substrate-bound protomers in the middle of the spiral are stable and bound to ATP (P2-P4) while the protomers towards the seam are in a post-hydrolysis ADP or apo state (Fig. 6a and Supplementary Fig. 6). This configuration agrees with the model for the clockwise rotary cycle of AAA+ translocases in which hydrolysis by the lowest protomer in the spiral (P1) triggers substrate release and ATP binding facilitates re-binding to the next substrate contact site^1,36^. The pore loop spacing along the substrate and conformational changes between the engaged and disengaged states are consistent with a two amino acid translocation step, similar to previous structures, but is smaller than step sizes reported for ClpA by single molecule^50^ and transient state kinetic methods^51^. This difference could be due to the use of ATPγS compared to ATP, which may reduce the step to a fundamental size based on the pore loop spacing.

The key discovery of our work here is that for processive cycles of unfolding, ClpA must rotate around the apical surface of ClpP in a regulated manner in order to maintain alignment of the spiral seam with the hexamer:heptamer mismatch (Fig. 6c and Supplementary Video 4). Additionally, while binding by IGF/L loops is well-understood to trigger gate-opening in ClpP, the conformational plasticity and asymmetric binding interactions we identify reveal new insight into how these loops facilitate allosteric regulation by ClpP^51,52^ and enable the AAA+ spiral to engage the ClpP planar surface during substrate translocation. A number of proteolysis machines operate as hexamer:heptamer assemblies^3,4^, thus a key question emerges as to how this mechanism is conserved. Indeed, recent structures of ClpXP reveal its IGF loops are similarly flexible, supporting a conserved rotary mechanism^56^. Notably, assembly of the eukaryotic Rpt and archaeal PAN AAA+ hexamer with its respective 20S core involves engagement by C-terminal HbYX motifs and gate-opening of the 20S^57,58^. For the 26S proteasome, alternating interaction by 3 of the asymmetric Rpt subunits is important for unfolding^59^ and a number of different engagement states have been identified^60^. Thus these interactions are similarly dynamic and a “wobbling” mechanism involving alternating HbYX interactions has been proposed for regulating gate opening^58^. While recent structures reveal a conserved spiral staircase arrangement of Rpt^40,61^ and PAN^62^, data supporting a rotation of the AAA+ hexamer around 20S, to our knowledge, has not been determined. Nonetheless, similar to our findings with ClpAP, the flexibility and distinct engagement states of the connecting HbYX motifs indicate conformational plasticity across the hexamer:heptamer interface is likely critical for coupling ATP hydrolysis and substrate unfolding with proteolysis. Furthermore, the symmetry mismatch may be critical in theses proteases for alignment of an empty binding pocket on the protease with the AAA+ spiral, thereby utilizing binding asymmetry to guide the rotary ATPase cycle and directionality of substrate translocation.

## Supplementary Information

### Methods

#### Purification and analysis of ClpA, ClpP and RepA(1-25)-GFP

ClpA and ClpP were purified as previously described^51,63^. RepA 1-25 protein was expressed with a C-terminal His6-tag construct from the pDS56/RBSII plasmid. Transformed BL21 cells were inoculated in LB media with 100 ug/mL Ampicillin and grown at 37°C to OD600nm = ~0.6–0.8. The cell culture was induced with 1 mM IPTG for ~4 h at 30°C. Cell pellet was resuspended in 40 mM HEPES pH 7.4, 2 mM β-mercaptoethanol, and 10% glycerol with protease inhibitors (EDTA-free) (Roche) and then lysed by sonication. Following centrifugation (16,000 × g, 20 min, 30°C), the supernatant was applied to a Ni-NTA column (GE Healthcare) followed by a gradient elution from 20 mM imidazole to 500 mM imidazole. Purity was verified by SDS-PAGE and fractions were combined and concentrated into a storage buffer (40 mM HEPES pH 7.4, 500 mM KCl, 20 mM MgCl2, 10% glycerol (v/v), and 2 mM β-mercaptoethanol).

The RepA(1-25)-GFP degradation assay (Supplementary Fig 1b) was performed in triplicate and consisted of 6 μM ClpA, 7 μM ClpP, 1 μM RepA(1-25)-GFP and 2 mM nucleotide incubated in buffer containing 50 mM Tris-HCl pH 7.5, 150 mM KCl, 10 mM MgCl_2_, and 1 mM DTT. Aliquots of the reaction were separated from the reaction at the specified time points and quenched in 2% SDS buffer, heated for 10 min and ran onto an acrylamide gel. The bands were visualized using silver staining (Sigma-Aldrich).

Size exclusion chromatography (SEC) analysis and purification was performed by incubating 36 μM ClpA, 42 μM ClpP, 30 μM RepA(1-25)-GFP and 2 mM ATPγS in 50 mM Tris-HCl pH 7.5, 150 mM KCl, 10 mM MgCl_2_, 1 mM DTT for 15 minutes. The complex incubation reaction was then injected onto a Superose 6 Increase 3.2/300 column (GE Healthcare) and the eluted peaks were analyzed using SDS-PAGE.

#### Cryo-EM Data Collection and Processing

The fraction corresponding to the largest molecular weight complex from SEC (Supplementary Fig. 1a) was isolated and incubated with 1 mM ATPγS before applying to glow discharged holey carbon grids (R 1.2/1.3; Quantifoil), plunge frozen using a vitrobot (Thermo Fischer Scientific) and imaged on a Titan Krios TEM (Thermo Fischer Scientific) operated at 300 keV and equipped with a Gatan BioQuantum imaging energy filter using a 20eV zero loss energy slit (Gatan Inc). Movies were acquired in super-resolution mode on a K3 direct electron detector (Gatan Inc.) at a calibrated magnification of 58,600X corresponding to a pixel size of 0.4265 Å/pixel. A defocus range of 1.2 to 2.0 µm was used with a total exposure time of 2 seconds fractionated into 0.2s subframes for a total dose of 68 e^−^/Å^2^ at a dose rate of 25 e^−^/pixel/s. Movies were subsequently corrected for drift using MotionCor2 (10.1038/nmeth.4193) and were Fourier-cropped by a factor of 2 to a final pixel size of 0.853 Å/pixel

A total of ~17,000 micrographs were collected over two different datasets. The two datasets were processed separately and then were combined at the end. All the data-processing was performed in cryosparc2^64^. For particle picking, templates were generated from 100 particles, in which only side-views were selected. After inspecting the particles picked, approximately 1.8 million particles were extracted. 2D classification was performed to remove contamination and junk particles, which amounted to ~44% of the dataset. A five-class ab-initio reconstruction was performed from the particle set, and was used for initial classification steps.

To identify different conformations and improve the resolution, heterogenous refinement was performed with 5 different classes (Supplementary Fig. 1j). Following this first round, maps showing high resolution features, which accounted for ~64% of the 512,000 particles going into 3D, were kept and grouped together. Another round of heterogenous refinement with 5 different classes was then performed. Following this second round, two unique states, ClpAP^Eng^ (19%, 169,000 particles) and ClpAP^Dis^ (33%, 97,000 particles), were identified. Particles associated with each class were combined and homogenous refinement was performed separately on each state. The same procedure was completed on the second dataset, which produced similar results in which ~52% of the 417,000 going into 3D were kept after the first round of classification followed by 34% in the Disengaged state and 17% in the Engaged State.

For the final refinement steps all the particles associated with either ClpAP^Eng^ (169,000 particles) or ClpAP^Dis^ (314,000 particles) were combined together and underwent Homogenous refinement. Due to the flexibility of the mobile protomers, Non-Uniform refinement was performed to improve the resolution. The final resolution of ClpAP^Eng^ was 3.1Å and ClpAP^Dis^ was 3.0Å (Supplementary Fig. 1d-h).

To better improve the resolution of the mobile protomers following Non-Uniform refinement, all the particles from both the ClpAP^Eng^ and ClpAP^Dis^ states underwent particle subtraction. Particle subtraction was performed in which the bottom half of ClpP was subtracted. A local-refinement was then performed, in which the fulcrum position was set to the center of ClpA. The same procedure was completed on the second state.

#### Molecular Modeling

An initial model for ClpA was obtain by using a ClpB structure (pdb 5ofo)^35^ and generated in SWISS-MODEL^65^ and the initial model for ClpP was taken directly from a ClpP crystal structure (pdb 1yg6)^46^ previously solved. Both initial models were docked into the EM maps using the UCSF chimera’s function *fit in map*^66^. Initial refinement was performed using Phenix^67^ with 1 round of simulated annealing and morphing and 5 rounds of real-space refinement that included minimization_global, rigid_body, adp, local_grid_search, secondary structural restraints and non-crystallographic symmetry (NCS) restraints. The resulting model then underwent real space refinement in Coot^68^. Nucleotides were added in manually using Coot and real space refinement using cif files generated for ADP and ATPγS in Phenix eLBOW^69^.

Density for the ClpA focus refinement was higher quality than the full map, therefore was used to model individual protomers using Rosetta Comparitive Modeling (RosettaCM)^70,71^. The structures for ClpA (pdb 1r6b)^72^, Hsp104 (pdb 5d4w and 5vjh)^36^, ClpB BAP form (pdb 5og1)^35^ and PTEX (pdb 6e10)^41^ were determined as homology models with HHpred^73^ and used to constrain model refinement in Rosetta CM with *template_weight=0* and the initial model with *template_weight=1*. The lowest energy models were examined by eye to ensure the model fit into the density, the protomer was placed into the context of the whole structure and the Rosetta Relax protocol was run on the full complex.

Rosetta Enumerative Sampling (Rosetta ES) was used to de novo build in the IGL loops and NT loops for each protomer^74^. The ClpA residues 612 to 628 were deleted from each protomer and Rosetta ES was run to rebuild the loops with a beamwidth of 32. The resulting model with rebuilt IGL loops was added into the full model and the Rosetta Relax protocol was run. Residues 16 to 32 from ClpP were deleted from each protomer and the same RosettaES parameters were used to build in the NT loops, followed by the Rosetta Relax protocol.

## Supplementary Figures

**Supplementary Figure 1:**
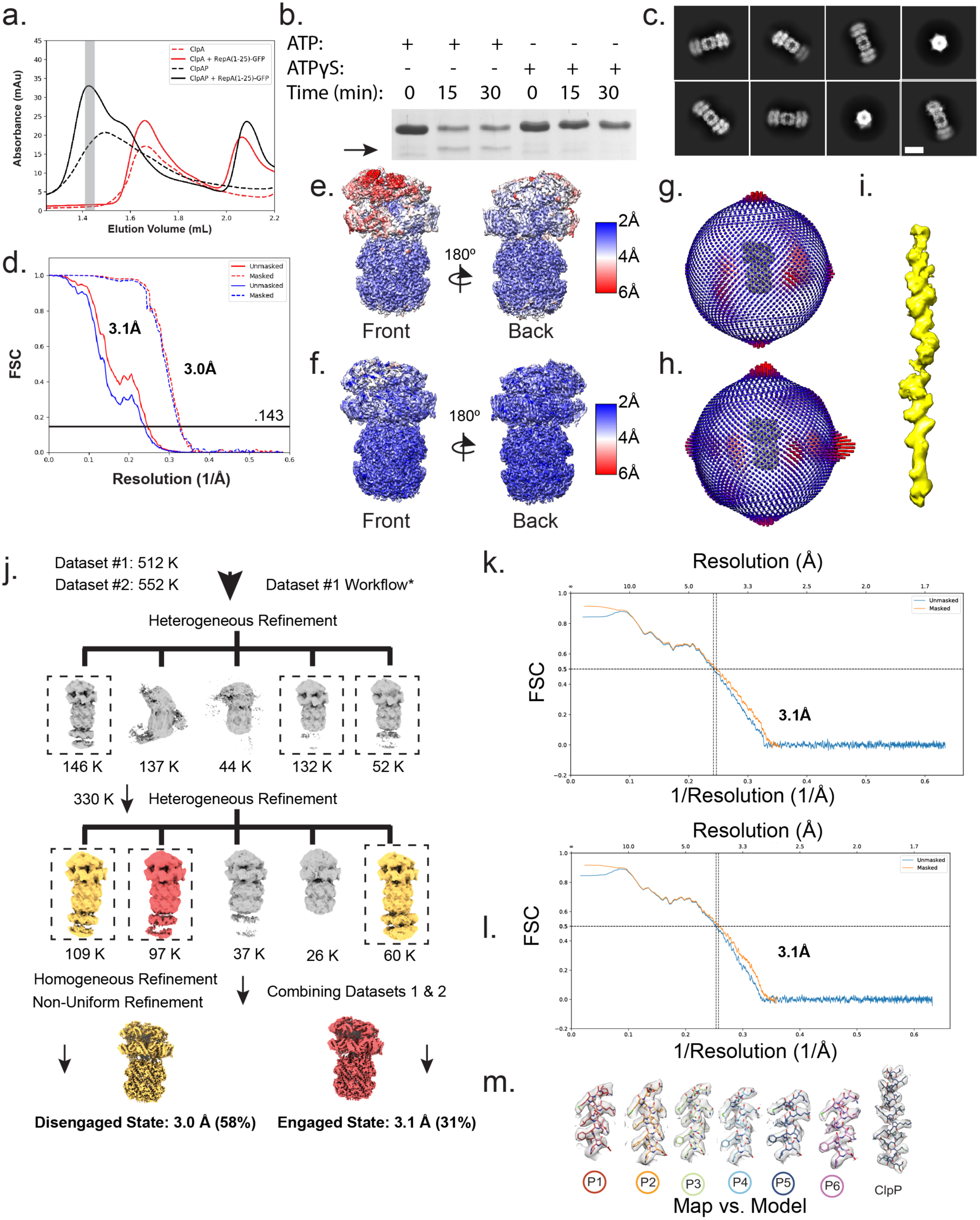
Analysis and cryo-EM Reconstruction of the ClpAP-RepA-GFP complex. **a,** Size exclusion chromatography (SEC) trace of the components and formed ClpAP complex following incubation with RepA^1–25^-GFP and ATPyS. Absorbance trace at 280 nm is shown for ClpA alone (red, dashed), ClpA with RepA^1–25^-GFP (red, solid), ClpAP alone (black, dashed) and ClpAP with RepA^1–25^-GFP (black, solid). **b,** RepA^1–25^-GFP degradation assay in the presence of ATP and ATPγS. Gel bands corresponding to RepA^1–25^-GFP are shown. **c,** Reference-free 2D class averages of ClpAP bound to RepA^1–25^-GFP. The scale bar equals 125 Å. **d,** Gold standard FSC-curves for the final refinement of both ClpAP^Eng^(blue) and ClpAP^Dis^(red) of the ClpAP-RepA(1-25)-GFP complex. The local resolution map and particle distribution plot of ClpAP^Eng^ (**e,g**) and ClpAP^Dis^ (**f,h**). **i,** Masked map of the substrate density for ClpAP^Dis^ at a reduced threshold showing continuous density spanning translocation channel. **j,** 3D classification scheme used to identify the two different states in the ClpAP-RepA^1–25^-GFP dataset. Dotted boxes represent the classes in which the particles were pooled together for further classification and refinement. The maps for ClpAP^Eng^ (red) and ClpAP^Dis^ (yellow) are colored accordingly. Map vs. Model FSC of ClpAP^Eng^ (**k**) and ClpAP^Dis^ (**l**) following atomic modeling in Rosetta. **m,** The EM density and atomic model of ClpAP-RepA^1–25^-GFP D2 helix D, which includes residues 571 to 583 for each protomer and ClpP helix E from a single protomer.

**Supplementary Figure 2:**
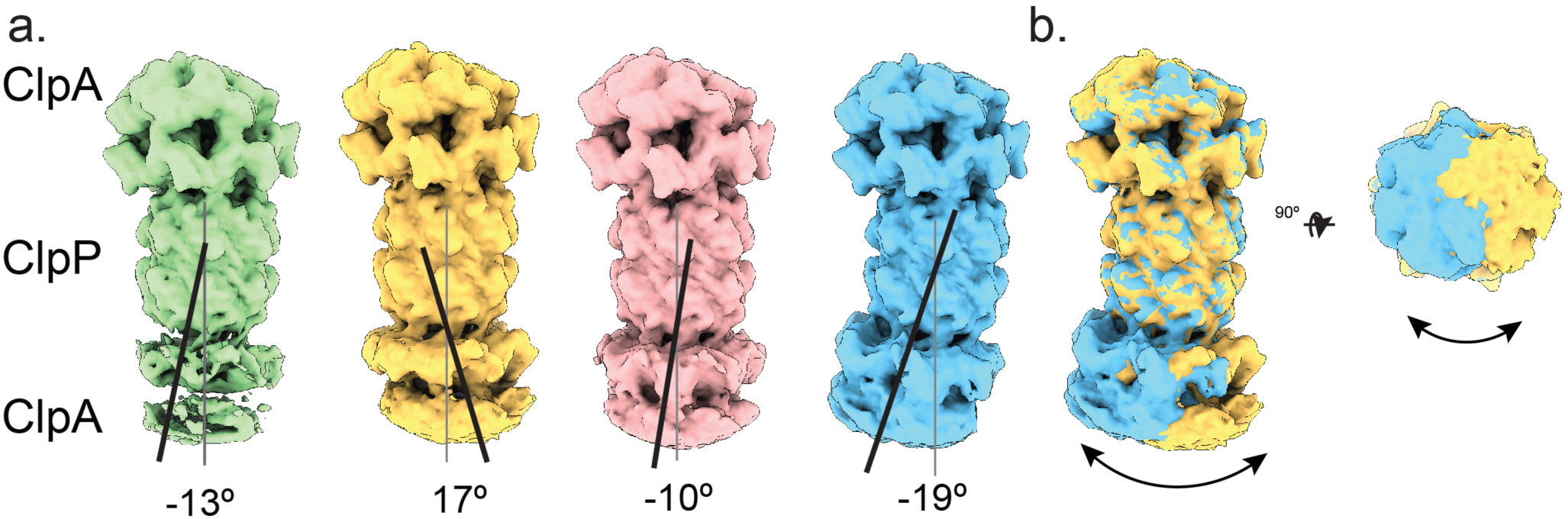
Pivot of ClpP around ClpA. **a,** Different classes of double-capped ClpAP identified by 3D classification with the position of ClpA substrate channel (black) and ClpP protease gate and chamber (grey) indicated. Maps are aligned to the upper ClpA hexamer in order to visualize rotations in the lower ClpA. The degree offset is indicated. **b,** Overlay of the two EM maps with the biggest offset of the channels, side view (left) and top view (right).

**Supplementary Figure 3:**
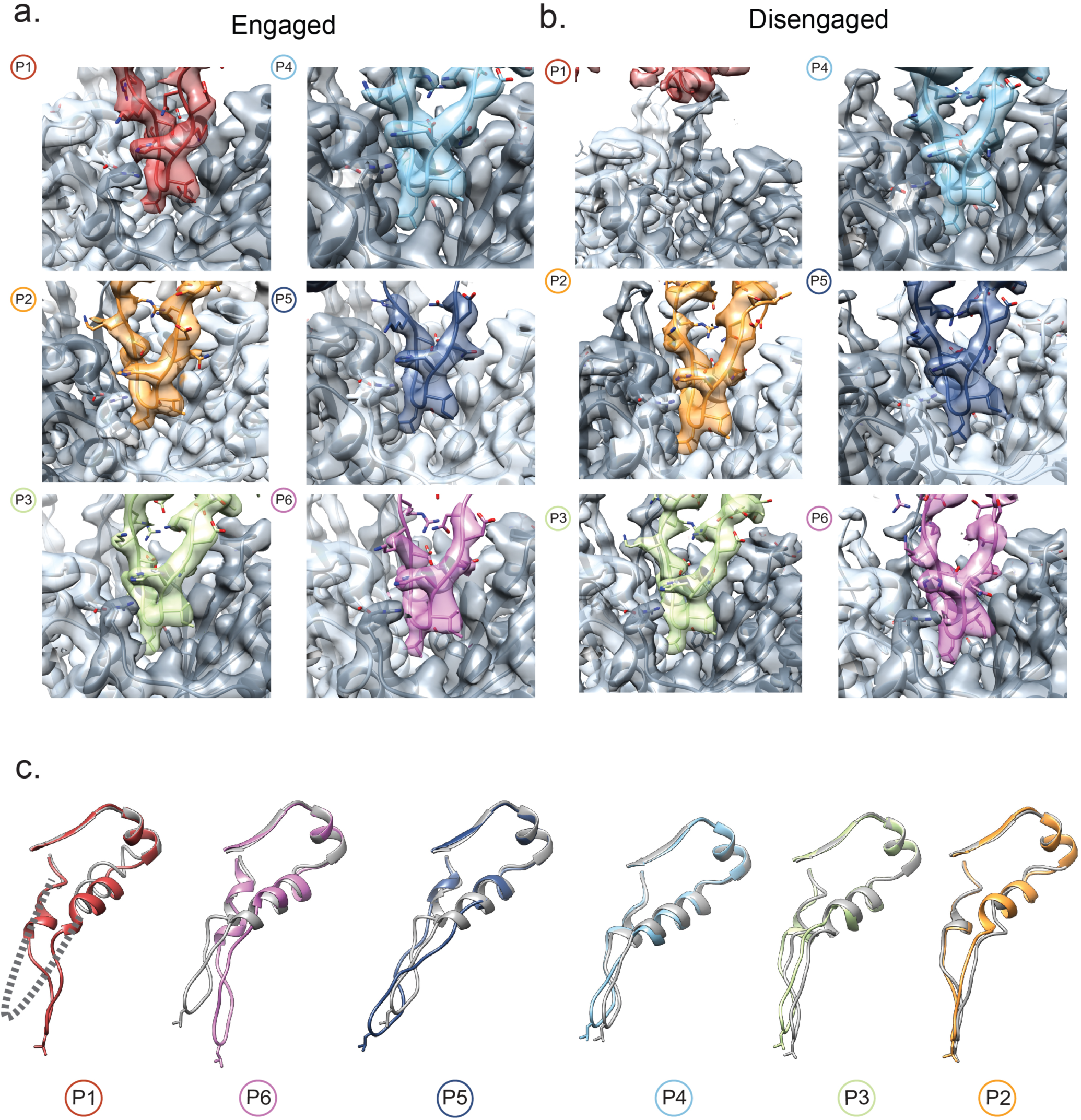
IGL-loop plasticity and comparison. EM map and model of the IGL-loops bound in the hydrophobic pocket for the engaged state (**a**) and disengaged state (b), protomer indicated by color and number. **c,** Overlay of IGL-loops of the engaged state (grey) and disengaged state (colored by protomer) laid out after alignment to the residues (638-649) above the IGL-loop. For protomer P1 in the disengaged state flexible loop residues that were not resolved are indicated (dashed line).

**Supplementary Figure 4:**
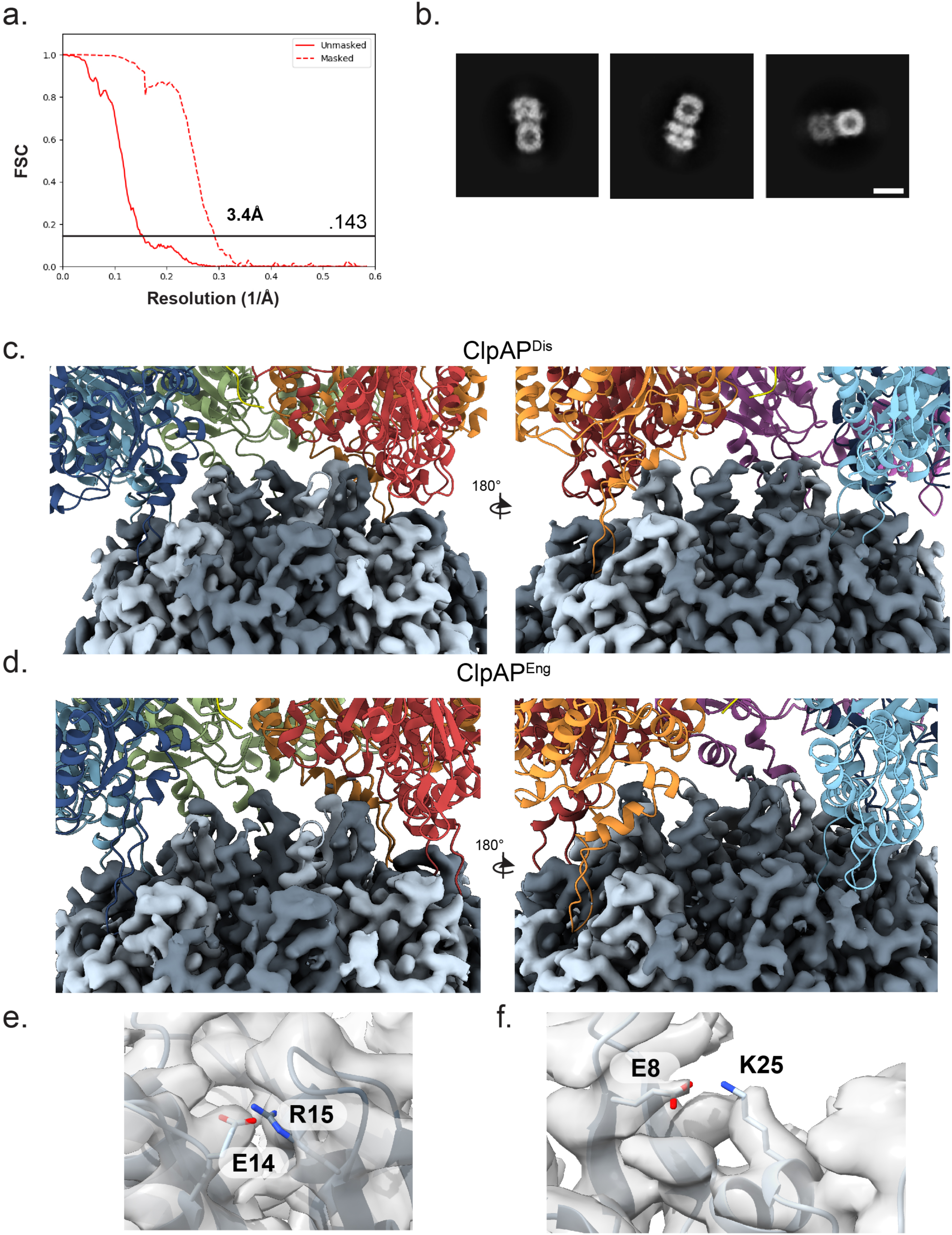
Single capped ClpAP structure and ClpP N-terminal loop interactions. **a,** Gold standard FSC curve of the single capped ClpAP structure. **b,** 2D reference free class averages of the single capped ClpAP structure. The scale bar equals 125 Å. The map vs model for ClpP N-terminal gating loops and the model for ClpA and substrate for ClpAP^Dis^(**c**) ClpAP^Eng^(**d**). Map vs model of ClpP residues E14 and R15 (**e**), and E8 and K25 (**f**).

**Supplementary Figure 5:**
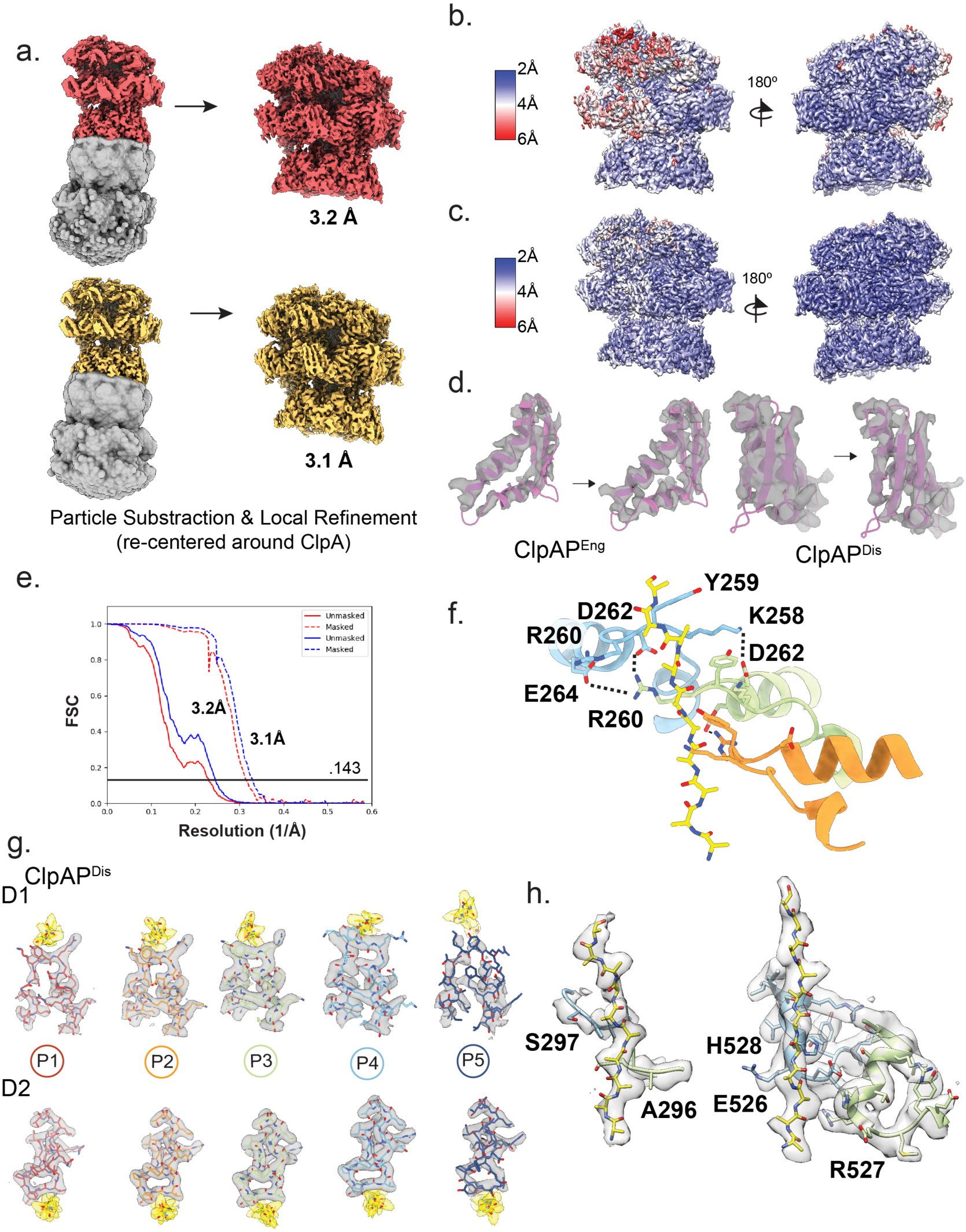
Particle Subtraction and Focus Refinement of ClpAP^Eng^ and ClpAP^Dis^. **a,** EM map with mask (grey) used for particle subtraction for both the engaged (top) and disengaged (bottom) states with the EM map after refinement (right). The local resolution map of ClpAP^Eng^ (**b**) and ClpAP^Dis^ (**c**). **d,** The EM density of different sections of P6 of both ClpAP^Eng^ (left) and ClpAP^Dis^ (right), before and after particle subtraction and focus refinement. **e** Gold standard FSC curve of both focus maps for ClpAP^Eng^ (red) and ClpAP^Dis^ (blue). **f,** Model of ClpAP^Dis^ (colored by protomer) with the D1 Tyr-containing pore loops residues interacting with adjacent pore loop residues. **g,** EM map and model of each Tyr-containing pore loop in ClpAP^Dis^ for both D1 (top) and D2 (bottom), the substrate channel density is colored yellow. **h,** Model of ClpAP^Dis^ (colored by protomer) with the Tyr-containing pore loops residues interacting with adjacent pore loop residues.

**Supplementary Figure 6:**
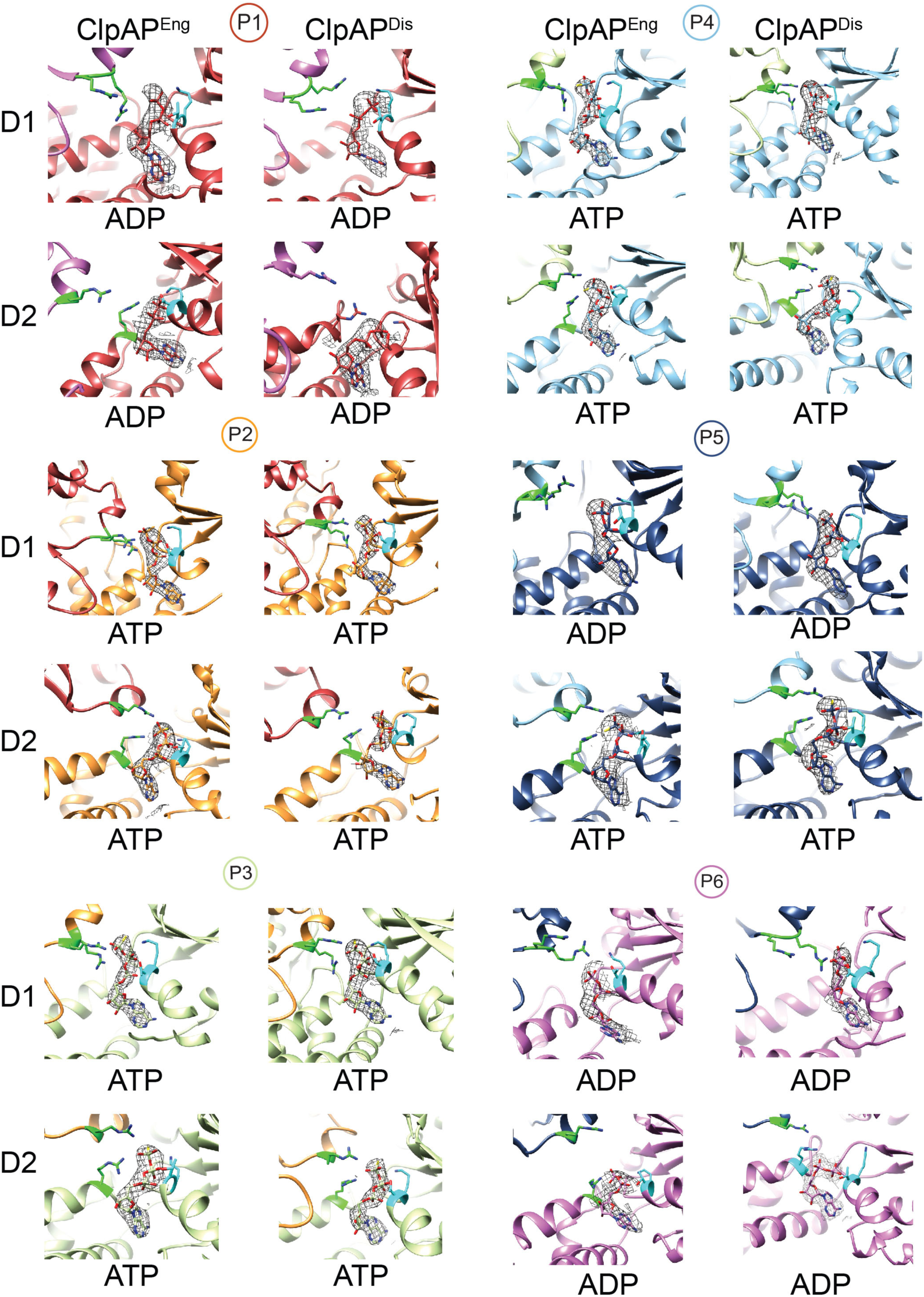
Nucleotide States of ClpAP^Eng^ and ClpAP^Dis^. EM density map of the nucleotide at each nucleotide pocket of the engaged and disengaged state of both D1 and D2.

---

**Supplementary Table 1.** Cryo-EM data collection, refinement and validation statistics.

Cryo-EM data collection, refinement and validation statistics
|  | Disengaged<br>State | Engaged<br>State | Focus<br>Disengaged<br>State | Focus<br>Engaged<br>State |
| --- | --- | --- | --- | --- |
|  | EMDB<br>20845<br>PDB<br>6UQE | EMDB<br>20851<br>PDB<br>6UQO | EMDB<br>20852<br>PDB<br>6UQP | EMDB<br>20848<br>PDB<br>6UQH |
| **Data collection and processing** |  |  |  |  |
| Microscope |  |  | Titan Krios |  |
| Camera |  |  | K3 |  |
| Magnification |  |  | 58,600 |  |
| Voltage (kV) |  |  | 300 |  |
| Data acquisition software |  |  | SerialEM |  |
| Exposure navigation |  |  | Image Shift t |  |
| Electron exposure (e-<br>/Å<sup>2</sup>) |  |  | 68 |  |
| Defocus range (μm) |  |  | 1.2-2 |  |
| Pixel size (Å) |  |  | .853 |  |
| Symmetry imposed | C1 | C1 | C1 | C1 |
| Initial particle images<br>(no.) | 1,800,000 | 1,800,000 | 1,800,000 | 1,800,000 |
| Final particle images<br>(no.) | 314,000 | 169,000 | 314,000 | 169,000 |
| Map resolution (Å) | 3.0 | 3.1 | 3.0 | 3.2 |
| FSC threshold | .143 | .143 | .143 | .143 |
| Map resolution range (Å) |  |  |  |  |
| **Refinement** |  |  |  |  |
| Model resolution (Å) | 3.1 | 3.1 |  |  |
| FSC threshold | .143 | .143 |  |  |
| Model resolution range<br>(Å) |  |  |  |  |
| Map sharpening *B* factor<br>(Å<sup>2</sup>) | -112.9 | -103.2 | -108.6 | -98.0 |
| Model composition |  |  |  |  |
| Non-hydrogen atom | 48,402 | 48,522 |  |  |
| Protein residues | 6,161 | 6,174 |  |  |
| Ligands | 12 | 12 |  |  |
| *B* factors (Å<sup>2</sup>) |  |  |  |  |
| Protein | 33.7 | 33.4 |  |  |
| Ligand | 44.1 | 20.0 |  |  |
| R.m.s. deviations |  |  |  |  |
| Bond lengths (Å) | .025 | .026 |  |  |
| Bond angles (°) | 1.82 | 1.84 |  |  |
| Validation |  |  |  |  |
| MolProbity score | 1.37 | 1.25 |  |  |
| Clashscore | 3.52 | 2.58 |  |  |
| Poor rotamers (%) | .37 | .08 |  |  |

**Supplementary Video 1:**

Pivot of ClpA on ClpP between the engaged and disengaged states. The IGL-loop plasticity between the engaged and disengaged states is also shown.

**Supplementary Video 2:**

The substrate contacts of the Tyr-containing pore loops in both the engaged and disengaged states.

**Supplementary Video 3:**

Movement of seam protomers (P1, P5 and P6) between the engaged and disengaged states. The movie depicts a morph between the engaged and disengaged states and the two models are aligned to the stationary protomers (P2, P3 and P4).

**Supplementary Video 4:**

Overall dynamic translocation mechanism of ClpAP involving substrate release and IGL-loop binding and rebinding at the spiral seam.

